# Ecotoxicological effects of silver nanoparticles in marine mussels

**DOI:** 10.1101/2021.07.19.452197

**Authors:** A. Calisi, C. Lorusso, J.A. Gallego-Urrea, M. Hassellöv, F. Dondero

## Abstract

In the marine bioindicator species *M. galloprovincialis* Lam we predicted toxicity and bioaccumulation of 5 nm alkane-coated and 50 nm uncoated silver nanoparticles (AgNPs) along with Ag^+^, as a function of the actual dose level. We generated a time persistence model of silver concentration in seawater and used the Area Under the Curve (AUC) as independent variable in hazard assessment. This approach allowed us to evaluate unbiased ecotoxicological endpoints for acute (survival) and chronic toxicity (byssal adhesion). Logistic regression analysis rendered LC50_96h_ values of 0.68 ± 0.08; 1.00 ± 0.20; 1.00 ± 0.42 mg h L^−1^ respectively for Ag^+^, 5 nm and 50 nm AgNP posing no evidence the silver form is a necessary variable to predict the survival outcome. By contrast, for byssal adhesion regression analysis revealed a much higher toxicological potential of Ag^+^ vs AgNPs, 0.0021 ± 0.0009; 0.053 ± 0.016; 0.021 (no computable error for 50 nm AgNP) mg h L^−1^, and undoubtedly confirmed a role of the silver form.

Bioaccumulation was higher for Ag^+^ > 5 nm AgNP > 50 nm AgNP reflecting a parallel with the preferential uptake route / target organ.

We, eventually, provided a full range of toxicological endpoints to derive risk quotients.

## Introduction

OECD emphasized four metal or metal oxide engineered nanoparticles with high interest due to their inherent properties. According to the Nanotechnology Product Database (https://product.statnano.com), silver nanoparticles (AgNPs) are, currently, found in 950 consumer products belonging to 15 different industrial sectors, with medicine, textile, cosmetics, home appliance, environment and construction dominating the scene. In addition, in the 2007-2017 decade nearly 5000 new applications were registered in different fields, such as consumer, medical, agricultural and industrial (Sim et al., 2018). AgNPs attracted increasing interest due to their unique physical, chemical and biological (antibacterial) properties compared to their macro-scaled counterparts (Chen et al., 2009; Singh et al., 2015). Moreover, as other metal and metal oxides, AgNP can be obtained through green chemistry, therefore are promising sustainable next generation materials (Beyene et al., 2017; Sharma et al., 2009). AgNPs have peculiar physicochemical properties: high electrical and thermal conductivity, chemical stability, catalytic activity and non-linear optical behavior (Krutyakov et al., 2008). The physicochemical characteristics of metal-based NPs are fundamental to determine their environmental fate as well as speciation in water. Dissolution, aggregation and sedimentation of AgNP are likewise crucial processes, in particular in the marine environments, able to affect their bioavailability and toxicological properties (Darlington et al., 2009; Markus et al., 2015). From an ecotoxicological point of view, the scientific community is still arguing if the environmental release of engineered nanoparticles may pose an ecological risk greater than that determined by bulk materials (non-nano) and/or the ionic forms. Several reports question for a harsher effect of nanomaterials in different model species (Shaw and Handy, 2011; Schultz et al., 2016; Bourdineaud et al., 2021; Parsal and Kumar 2021), however, exceptions were reported that need consideration (an example is given in Lahive et., 2017). With regards to the marine environment despite the information has linearly increased in the last decade, the big picture of NP toxicity is still limited due to additional technical difficulties. Indeed, the behavior of engineered NPs in the seawater is highly influenced by aggregation, agglomeration sedimentation, mobility, speciation and dissolution (Timerbaev et al., 2021), confounding factors that increase the uncertainty in hazard and risk assessment.

Several studies in the marine matrix focused on primary producers and consumers, such as algae and bivalves i.e. *Mytilus* congeners. Mussels are - indeed-widely recognized as a NP sensitive species with also an ecological relevance (Baun et al., 2008; Moore, 2006; Canesi et al., 2012). A quick literature survey says that great attention was given to AgNPs with different coatings (polyvinyl pyrrolidone dominated the scene) and sizes (from 5 to 100 nm are well represented), but paradoxically, the vast majority of the studies on mussels focused on sublethal effects viz. biomarkers. Moreover, most reports were centered on a few selected concentrations (the mode is in the 0.010-0.10 mg L^−1^ bin). To our knowledge, in fact, if excluding Katsumiti et al. (2015) provding cumulative toxicity curves along with LC50 for hemocytes and isolated gills cells exposed to maltose AgNP of different sizes; and Auguste et al. (2018) providing cumulative toxicity curves along with EC50 for hemocytes lysosomal membrane stability (LMS) and larval development, no full range effects for acute and chronic toxicity has been yet determined in adult specimens of the *Mytilus* congeners. By contrast, a few dozen studies have reported biomarker data.

The aim of this work, therefore, was to fill this gap providing a comprehensive acute (survival) and chronic (byssal adhesion) hazard assessment in *M. galloprovincialis* Lam. To this aim, 5 nm alkane-coated and 50 nm uncoated AgNPs were used in a nominal concentration range comprising 5 magnitude orders (0.001 – 10.0 mg L^−1^). To establish the genuine toxicological potential of AgNPs and allow an unbiased comparison with ionic silver (Ag^+^), we first generated a persistence model to calculate the variation of the silver titer as a function of time in seawater and, hence, we used the Area Under the Curve (AUC), i.e. the silver integrated dose, as the principal driver of toxicity and bioaccumulation in (multiple) regression analysis.

## Materials and Methods

### Materials

Ionic silver in the form of nitrate was obtained from Sigma-Aldrich (99% purity). 5nm alkane-coated AgNP (particle range 3–8 nm) were obtained from Amepox sp. Z o. O. (Poland) in the form of a stable aqueous 1 g L^−1^ dispersion. According to the manufacturer these particles have an alkane-coating accounting for 18% weight whose composition represents an industrial secret and therefore its exact composition is unknown. Primary characterization of Amepox AgNP was previously reported (Ribeiro et al., 2014). A summary of primary characterization is presented in Section 3.1 along with secondary characterization.

50 nm uncoated AgNP were obtained from NanoTrade s.r.o. (Czech Republic) in the form of a grey / brownish powder. Primary characterization of NanoTrade AgNP was previously reported (Diez-Ortiz et al., 2015). Briefly, TEM analysis showed primary particles in the range of 50-80 nm in size, forming larger nano-to micron-sized aggregates.

### Nanoparticle secondary characterization and aggregation experiments

Characterization of the stock solutions were done by dynamic light scattering. For the aggregation experiments the instrument used was a Zetasizer (Malvern Panalytical Ltd, Malvern, UK), with the Zetasizer software version 6.20.

Different seawater media was used to test for nanoparticle stability: 35‰ Artificial Sea Water (ASW; Instant Ocean, VA, USA), 35‰ natural sea water (NSW) and 35‰ Natural Sea water filtered (NSWf) with very similar results.

Short term experiments were performed to investigate initial aggregation rates of the 5 nm AgNPs. The stock solutions were prepared before the experiments and mixed with the media and MQ to the desired concentrations in regular DLS cuvettes, mixed briefly using a vortex mixer and inserted immediately in the instrument. The measurement was started at a fixed attenuator and measurement position with the correlation time set to 2 seconds and 120 data points were generally obtained.

During long term experiments (up to 4 days) the first measurement (day zero) was obtained by creating an average result from the short term data points. The cuvettes were stored at dark and three measurements were performed (3 runs of 20 seconds each) in the following days. To evaluate the effect of particles sedimentation, the samples were shaken after performing the measurement and a new measurement was done.

### Exposure experimental design

Adult mussels (*Mytilus galloprovincialis* Lam.) were sampled from a natural population in front of Gabicce Mare (Italy, North Adriatic Sea) and transported to laboratory in a cool box under controlled condition (0-4°C). Animals (n= 600) were selected to form homogeneous groups in size (5-6 cm), separated from one another by carefully cutting off the byssal threads and washed with\ seawater. Then, mussels were acclimatized for at least 4 weeks in static tanks containing filtered 35‰ aerated artificial sea water ASW (Instant Ocean, VA, USA), pO_2_ > 8.0 mg/L, pH 8.1 ± 0.5. Temperature was kept constant at 16±1°C and animals fed daily during acclimatation with a commercial nutritive solution for marine invertebrates (Marine Liquifry, Interpret, UK).

Mussels were randomly assigned to experimental (Ag treated) and reference not-exposed groups. Five different nominal exposure levels –from 10 mg L^−1^ to 0.001 mg L^−1^ Ag with a log10 series, and 2 reference control groups were set-up for each replicated experiment (n=4). One control group contained the alkane-coating at 1.8 mg L^−1^ to resemble the worst-case condition, howeer, control data were merged during the analysis since no significant effects were found due to coating. The nominal concentrations of 50 nm AgNP were adjusted to obtain similar values of the 5 nm AgNP that account for the alkane coating (see paragraph Table 1).

**Table1.**
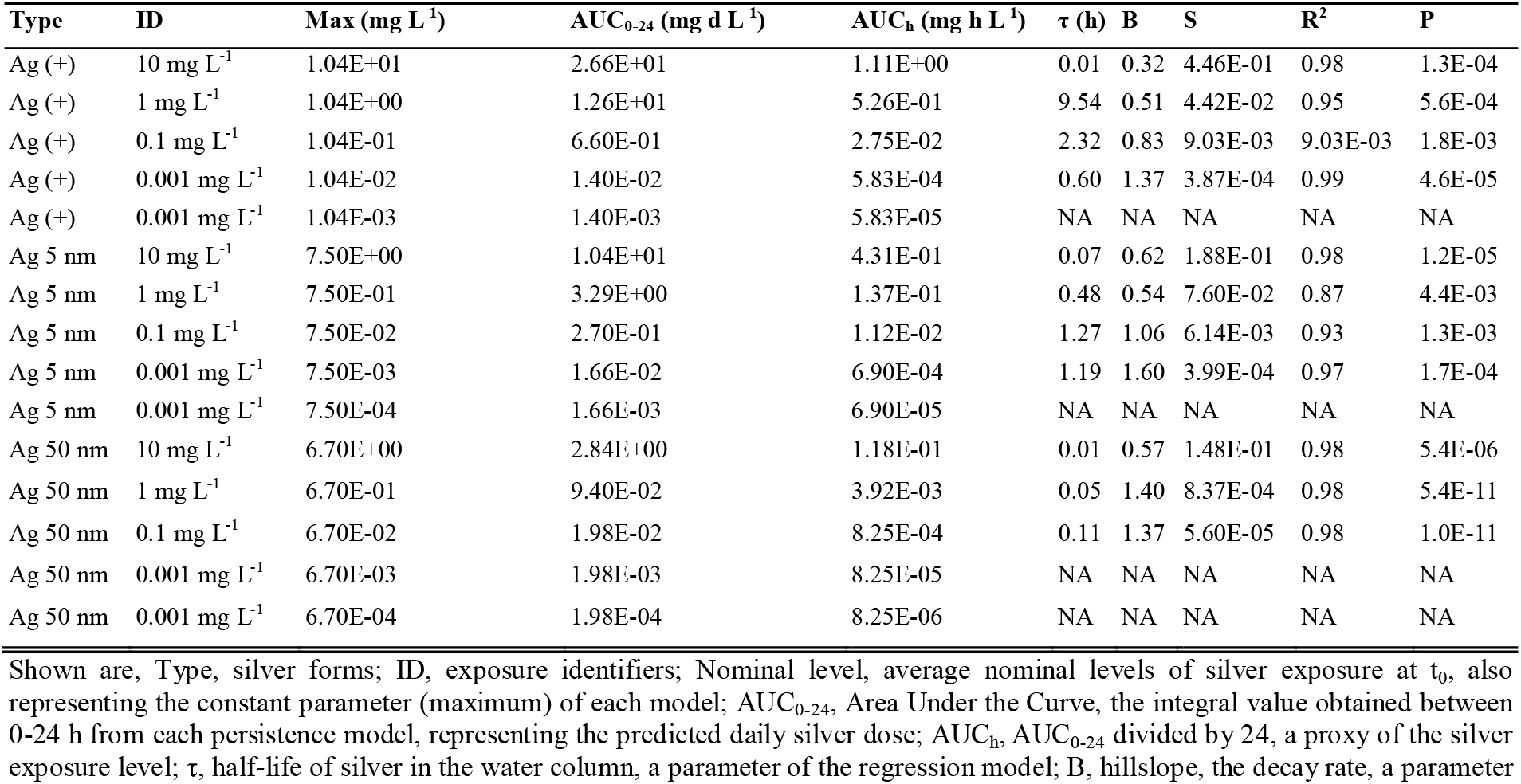
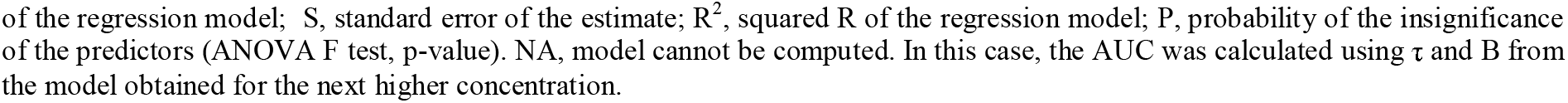
Actual silver doses and persistence model parameters (logistic regression)

| Type | ID | Max ($\text{mg L}^{-1}$) | $\text{AUC}_{0-24}$ ($\text{mg d L}^{-1}$) | $\text{AUC}_h$ ($\text{mg h L}^{-1}$) | $\tau$ (h) | B | S | $R^2$ | P |
| --- | --- | --- | --- | --- | --- | --- | --- | --- | --- |
| Ag (+) | 10 $\text{mg L}^{-1}$ | 1.04E+01 | 2.66E+01 | 1.11E+00 | 0.01 | 0.32 | 4.46E-01 | 0.98 | 1.3E-04 |
| Ag (+) | 1 $\text{mg L}^{-1}$ | 1.04E+00 | 1.26E+01 | 5.26E-01 | 9.54 | 0.51 | 4.42E-02 | 0.95 | 5.6E-04 |
| Ag (+) | 0.1 $\text{mg L}^{-1}$ | 1.04E-01 | 6.60E-01 | 2.75E-02 | 2.32 | 0.83 | 9.03E-03 | 9.03E-03 | 1.8E-03 |
| Ag (+) | 0.001 $\text{mg L}^{-1}$ | 1.04E-02 | 1.40E-02 | 5.83E-04 | 0.60 | 1.37 | 3.87E-04 | 0.99 | 4.6E-05 |
| Ag (+) | 0.001 $\text{mg L}^{-1}$ | 1.04E-03 | 1.40E-03 | 5.83E-05 | NA | NA | NA | NA | NA |
| Ag 5 nm | 10 $\text{mg L}^{-1}$ | 7.50E+00 | 1.04E+01 | 4.31E-01 | 0.07 | 0.62 | 1.88E-01 | 0.98 | 1.2E-05 |
| Ag 5 nm | 1 $\text{mg L}^{-1}$ | 7.50E-01 | 3.29E+00 | 1.37E-01 | 0.48 | 0.54 | 7.60E-02 | 0.87 | 4.4E-03 |
| Ag 5 nm | 0.1 $\text{mg L}^{-1}$ | 7.50E-02 | 2.70E-01 | 1.12E-02 | 1.27 | 1.06 | 6.14E-03 | 0.93 | 1.3E-03 |
| Ag 5 nm | 0.001 $\text{mg L}^{-1}$ | 7.50E-03 | 1.66E-02 | 6.90E-04 | 1.19 | 1.60 | 3.99E-04 | 0.97 | 1.7E-04 |
| Ag 5 nm | 0.001 $\text{mg L}^{-1}$ | 7.50E-04 | 1.66E-03 | 6.90E-05 | NA | NA | NA | NA | NA |
| Ag 50 nm | 10 $\text{mg L}^{-1}$ | 6.70E+00 | 2.84E+00 | 1.18E-01 | 0.01 | 0.57 | 1.48E-01 | 0.98 | 5.4E-06 |
| Ag 50 nm | 1 $\text{mg L}^{-1}$ | 6.70E-01 | 9.40E-02 | 3.92E-03 | 0.05 | 1.40 | 8.37E-04 | 0.98 | 5.4E-11 |
| Ag 50 nm | 0.1 $\text{mg L}^{-1}$ | 6.70E-02 | 1.98E-02 | 8.25E-04 | 0.11 | 1.37 | 5.60E-05 | 0.98 | 1.0E-11 |
| Ag 50 nm | 0.001 $\text{mg L}^{-1}$ | 6.70E-03 | 1.98E-03 | 8.25E-05 | NA | NA | NA | NA | NA |
| Ag 50 nm | 0.001 $\text{mg L}^{-1}$ | 6.70E-04 | 1.98E-04 | 8.25E-06 | NA | NA | NA | NA | NA |

Shown are, Type, silver forms; ID, exposure identifiers; Nominal level, average nominal levels of silver exposure at $t_0$ , also representing the constant parameter (maximum) of each model; $\text{AUC}_{0-24}$ , Area Under the Curve, the integral value obtained between 0-24 h from each persistence model, representing the predicted daily silver dose; $\text{AUC}_h$ , $\text{AUC}_{0-24}$ divided by 24, a proxy of the silver exposure level; $\tau$ , half-life of silver in the water column, a parameter of the regression model; $B$ , hillslope, the decay rate, a parameter
of the regression model; S, standard error of the estimate; $R^2$ , squared R of the regression model; P, probability of the insignificance of the predictors (ANOVA F test, p-value). NA, model cannot be computed. In this case, the AUC was calculated using $\tau$ and B from the model obtained for the next higher concentration.

Exposed mussels were treated for 24 or 96 h with each silver form, according to the test to be performed. Silver was added daily-along with water renewal-from freshly prepared stock water-suspension/solution or from the 5 nm stable stock solution. In general, 15 mussels per each condition/ replicate were tested. Animals were not fed during the exposures. After the acclimation, 15 mussels were randomly placed into new 21 L polypropylene plastic vessels with 35‰ aerated (110 L/h) ASW, pH 8.1 ± 0.5, at a density of 1 animal/L at 16°C. The 50 nm AgENPs stock solution was prepared according to the “PROSPEcT 2010: Protocol for Nanoparticle Dispersion” (PROSPEcT, 2010) (https://nanotechia.org/sites/default/files/files/PROSPECT_Dispersion_Protocol.pdf). Briefly, nanoparticles were weight and placed in a glass vial; a few drops of MilliQ water were added to create a concentrated nanoparticle paste and then more water was added to make up the desired concentration (0.5 – 1.0 g L^−1^). After sonication for 30 sec with an ultrasound probe at middle power intensity the suspension was stirred by hand for 10 s. Silver nitrate was prepared in ultrapure water at 0.5 g L^−1^. The silver titer was evaluated *ex post* in all stock solutions by chemical analysis (see below). Physical/chemical parameters (pH, T, O2) were checked at least daily along with ecotoxicological parameters.

### 96-h acute toxicity test – survival

Mortality was evaluated daily before the water renewal and at the end of the exposure, 96 h. Mussels were considered dead (and withdrew from the aquaria) when the lack of the adductor muscle activity was recorded. At least 4 independent exposure trials were carried out.

### 96-h chronic toxicity test – byssal adhesion

A simple binomial procedure was set up to test byssal performance in immersed mussels after 24 h exposure to Ag. Mussels were considered positive for adhesion if functional byssal threads of a specimen were found attached to the tank or to another individual mussel. At least 4 independent exposures per each condition were carried out.

### Determination of silver concentration in water samples

50 ml seawater samples were withdrawn from the exposure vessels at regular intervals of 1, 4, 24 h. For soluble silver, the water samples were ultracentrifuged at 100,000 g for 2 h at 20°C. Silver amounts at time zero (t_0_) were calculated from the stock solution concentration. Samples were analyzed for silver content by Inductively Coupled Plasma Mass-Spectrometry (ICP-MS) according to the unified referenced procedure UNI EN ISO 17294-2:2016 including certified materials as a reference standard. AgNP samples were acid digested before quantification according to the procedure EPA 3051A. The LOD for silver was 0.0005 mg L^−1^.

### Determination of silver concentration in mussel tissues

Total silver burdens were evaluated in whole soft tissues and isolated tissues, viz. gills and digestive glands from mussels exposed for 4 days to the different silver forms. Tissue biopsies were washed under cold tap water for 30 seconds, rinsed for 20 seconds in several volumes of cold 1 mM cysteine solution, rinsed twice in seawater, once in ultrapure water, damped and flash frozen in liquid nitrogen. Samples were stored at −80°C until needed. For metal content analysis, the soft tissue from single specimens were thawed and homogenized with the addition of 5 ml of ultrapure water per gram of tissue. Samples were acid digested according to the procedure EPA 3051A using a microwave oven by the addition of 10 volume of a 3:1 mixture of concentrated HCl:HNO_3_ (aqua regia) and further analyzed by ICP-MS analysis according to the referenced procedure UNI EN ISO 17294-2:2016 including certified materials as a reference standard. The limit of detection (LOD) was 0.005 mg kg^−1^

### Regression analysis and statistics

For the persistence model, non-linear regression of silver data in the water column was performed in MS-Excl (Microsoft Inc.) by parameter iteration using the three-parameter logistic function (3PL) and minimizing the least square residuals, essentially as described in Haanstra et al., 1985. Toxicity data was fit with the three-parameter logistic function (3PL) using SigmaPlot version 12.0, (Systat Software, Inc., San Jose California USA). Multiple logit regression was carried out using Stata version 14.2 (StataCorp, College Station, Texas USA).

Bioaccumulation data were log transformed and fit with (multiple) linear regression using SigmaPlot version 12.0 after checking for Residual Normality (Shapiro-Wilk test p > 0.05) and Constant Variance (Levene’s means test p > 0.05).

Differences in bioaccumulation patters between gills and digestive glands were tested by means of 1-way ANOVA (p <0.01) using Systat version 12.02 (Systat Software, Inc., San Jose California USA).

## 3. Results

### 3.1 Characterization of AgNP

Primary characterization (ultrapure water) of 5 nm AgNPs was performed elsewhere (Ribeiro et al., 2014) showing tiny particles ranging in size from 3 to 8 nm and forming aggregates wrapped in an organic layer (according to the manufacturer paraffin is used to protect particles and accounts for 18% in weight) which exhibits Z-average values (measured by dynamic light scattering) ranging from 60 to 130 nm when diluted to 10 mg L^−1^ in MQ water. Despite a stable behavior in pure water, the 5 nm AgNPs aggregated rapidly in seawater with faster kinetics at increasing concentrations (Figure 1, Panel A). In longer times, 24 h and plus, 5 nm AgNP tends to form micron-sized aggregates (Figure 1, Panel B-D).

**Figure 1.**
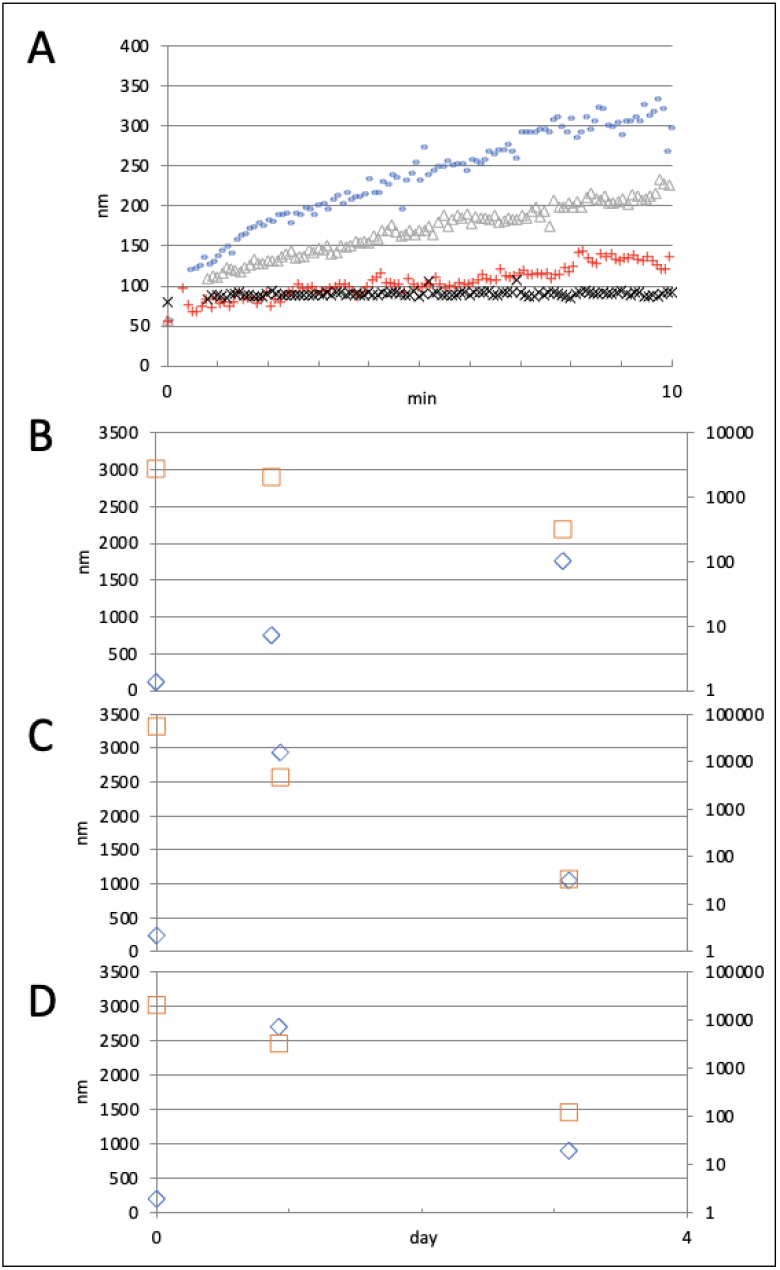
Pattern of 5 nm AgNP aggregation in seawater. Dynamic Light Scattering (DLS) analysis showed a consistent aggregation of 5 nm AgNPs with increasing hydrodynamic diameter (Z-average, nm) across time. Panel A, sudden aggregation in ASW at nominal concentrations of 5 nm AgNP (+, 1 mg L^−1^; Δ
, 5 mg L^−1^; –, 10 mg L^−1^; X, ultrapure water). Panel B-D, effect of longer time (day) on aggregate size at 1, 5 and 10 mg L^−1^. Micron sized AgNP aggregates appeared within 24 h (the renewal time in semi-static exposure) and showed to be unstable in seawater, as depicted by the count rate pattern. Legend, ⍰, Z-average, nm; ⍰, Derived count rate, kilo counts s^−1^. The phenomenon was time and concentration dependent. 50 nm AgNP could not be analysed by DLS due to an instantaneous precipitation in seawater.

DLS analysis in seawater was unreliable for the 50 nm AgNP due to instantaneous aggregation and precipitation that at 1-10 mg L^−1^ was clearly visible to the naked eye (data not shown). Indeed, large 3–5-micron sized granules could be observed in SEM analysis of mussel gills sampled at the end of the exposure (96 h) at different nominal exposure level (0.1-10 mg L^−1^). XRS analysis confirmed a high amount of Ag in aggregates found juxtaposed to or trapped into mussel gills suggesting these granules arose from primary 50 nm particles (Figure 2). Similar results but slightly smaller granules were obtained with 5 nm AgNP (data not shown).

**Figure 2.**
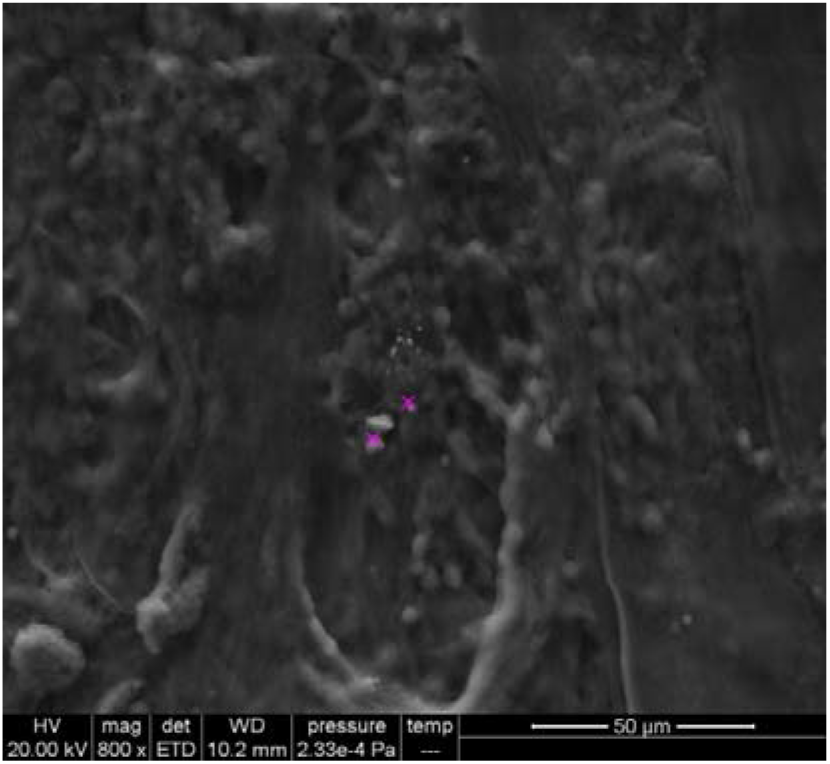
Scanning Electron Microscopy imaging of 50 nm AgNPs aggregates. A representative SEM photomicrograph depicts micron-sized aggregates (3-5 micron) arising from 50 nm Ag primary particles (white arrows). Granules were found juxtaposed to ethanol fixed gills sampled from mussels after 96 h semi-static exposure at 0.1 mg L^−1^ d (nominal level). Inlet panel, XRS analysis showing high Ag content in the granules (purple cross). Similar results were obtained for 5 nm Ag ENPs (data not shown).

### 3.2 Modeling of silver persistence in seawater

The variation of (total) silver concentration was measured in the water column during mussel exposure to the two particulate forms of silver −5 nm AgNP or 50 nm AgNP- or ionic silver (administered in the form of nitrate). Water was sampled at 1, 4 and 24 h from the beginning of each exposure. The titer at t_0_ was evaluated from the stock solution concentration. Five nominal exposure levels were used according to a log_10_ series. For the ionic form the nominal concentrations ranged from 10.4 to 0.0014 mg L^−1^. For the 5 nm AgNP, the stable stock solution obtained from the manufacturer showed a titer of 0.75 ± 0.45 g L^−1^ due to the alkane-coating, therefore nominal exposure levels started from 7.5 mg L^−1^. For 50 nm AgNP we, therefore, adjusted the concentration of the stock solutions to a similar level of the 5 nm AgNP. On average the stock solutions showed a titer of 0.67 ± 0.40 g L^−1^ therefore nominal exposure levels ranged from 6.7 mg L^−1^. For the sake of clarity all along the paper we used the 10-0.001 mg L^−1^ nominal levels of nitrate as labels to identify / compare the different exposures, also for 5 and 50 nm AgNPs. Table 1 reports details about the exposures and their identifiers.

Figure 3 shows the persistence of total silver in seawater with the severest drop for the 50 nm AgNP.

**Figure 3.**
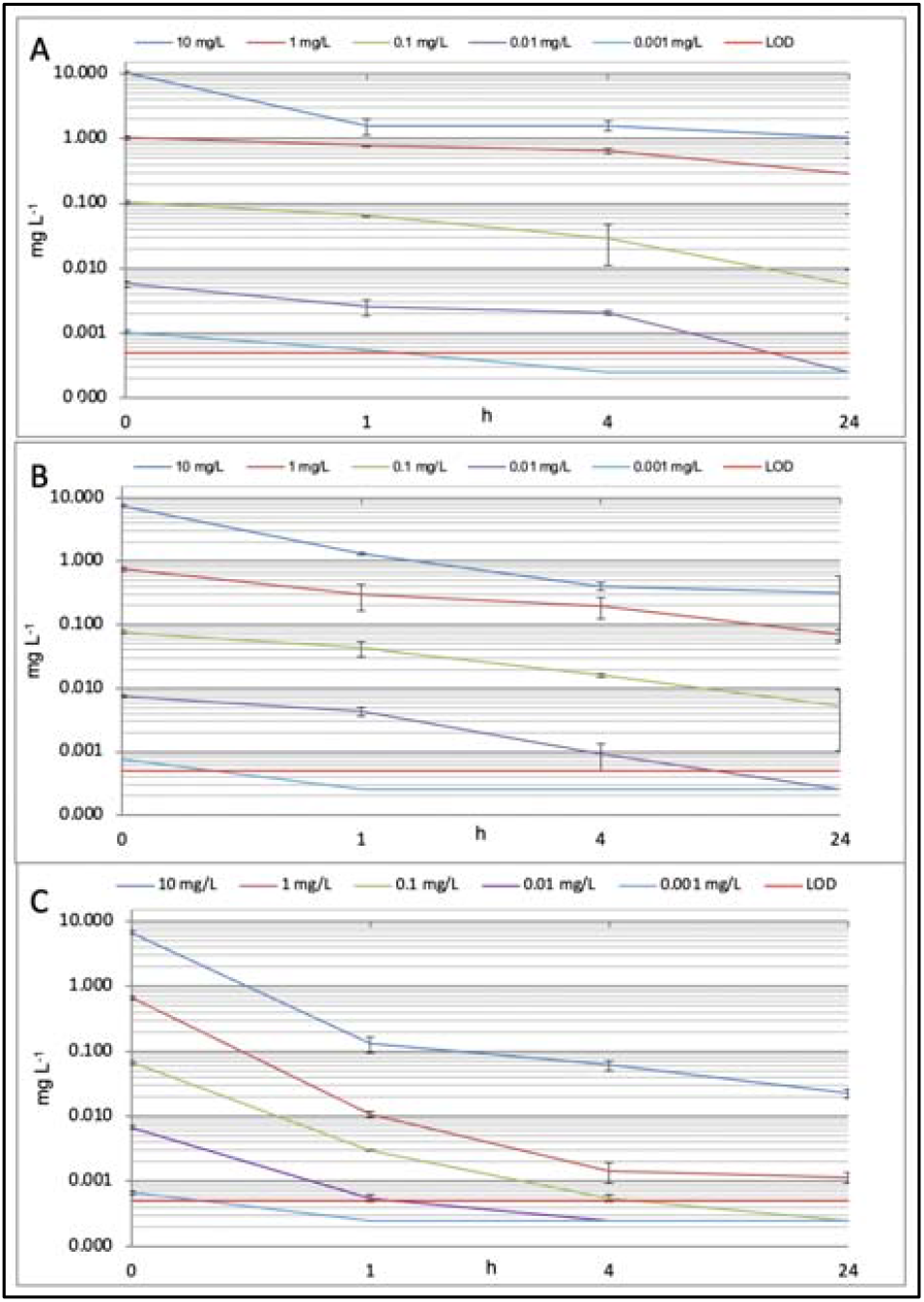
Persistence of Ag in seawater. Total Ag concentration was measured in the water column across the 24 h renewal time for each nominal exposure level. Concentrations at t_0_ were calculated from the titration of the stock solutions. Empirical Ag amounts were used to derive the integrated actual silver dose reported in Table 1. Shown are mean values +-SD (n=4). The horizontal red line represents the LOD level for silver, 0.0005 mg L^−1^. In these graphs and in modelling, samples below LOD were assumed equal to 0.00025 mg L^−1^. Legend. Panel A, Ionic silver; Panel B, 5 nm AgNP; Panel C, 50 nm AgNP.

Then, for each exposure conditions, we used a sigmoidal function to predict the silver concentration (y) in water using time (t) as the independent variable:

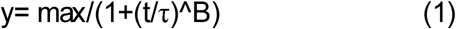

where **y** represents the silver concentration (mg L^−1^), t is time (h), τ is a model parameter representing the half-life of silver in the water column, B is a model parameter representing the hillslope of the curve, Max is a model parameter (constant) representing the silver concentration at t_0._ τ and B were parametrized to fit data into each model according to the least square residual using a spreadsheet software (Table 1). The persistence models allowed a high accuracy prediction of the the actual silver dose mussels were exposed to as the Area Under the Curve (AUC), i.e. the integral of the logistic function in 0-24 h (24 h is the renewal time). Table 1 reports the predicted AUC dosing per day (AUC_0-24_) and per hour (AUC_h_) - the latter representing a proxy of the actual dose level that can be useful for comparison with other works and external ecotoxicological datasets-along with the model coefficients (Max, τ and B) and the outcome of the ANOVA F test. All models for which a fair amount of data was available could be computed with a relatively low standard error of the estimate (S) and therefore were highly significant.

#### 96-h acute toxicity test – mortality in water

Survival rate was evaluated after 96 h exposure to 5 or 50 nm AgNPs or ionic silver at the five dose levels depicted in Table 1. Figure 4 shows a three parameters logistic regression model of survival vs the predicted (AUC_h_) dose for each silver forms. All models were highly significant according to ANOVA (p < 0.001) and could predict toxicity with a low standard error of the estimate (S), ranging from 3 to 6% (Table 2). Predicted toxicological endpoints (EC values) are presented in Table 2 along with the model parameters. EC50 for ionic Ag was apparently lower than the values obtained for the two nanoparticles 0.68 vs 1.0 AUC_h_ (mg h L^−1^). To test the significance of such difference and the possible dependence of survival on the silver form, we performed multiple logistic regression analysis on the whole survival dataset (without control data) including dummy variables for the classification of the *type of silver*. In this model, however, the type of silver variables did not appear necessary to predict the survival outcome (p = 0.711 for 5 nm AgNP; p = 0.242 for 50 nm AgNP).

**Table 2.** Logistic regression of silver toxicity: endpoints and model coefficients.

| Type | Effect | EC1 | EC5 | EC10 | EC20 | EC50 | SE | P | B | Max (%) | S | R<sup>2</sup> |
| --- | --- | --- | --- | --- | --- | --- | --- | --- | --- | --- | --- | --- |
| Ag (+) | survival | 3.20E-03 | 2.20E-02 | 5.30E-02 | 1.40E-01 | 6.80E-01 | 8.00E-02 | <0.0001 | 0.88 | 99.4 | 5.88 | 0.94 |
| Ag 5 nm | survival | 4.20E-03 | 3.00E-02 | 7.30E-02 | 1.90E-01 | 1.00E+00 | 2.00E-01 | <0.0001 | 0.88 | 99.3 | 3.43 | 0.91 |
| Ag 50 nm | survival | 3.90E-03 | 2.90E-02 | 7.10E-02 | 1.90E-01 | 1.00E+00 | 4.20E-01 | 0.02 | 0.92 | 97.7 | 4.17 | 0.70 |
| Ag (+) | adhesion | 1.96E-05 | 1.05E-04 | 2.24E-04 | 5.12E-04 | 2.10E-03 | 9.00E-04 | 0.013 | 1.17 | 85.3 | 7.64 | 0.96 |
| Ag 5 nm | adhesion | 2.53E-03 | 7.54E-03 | 1.24E-02 | 2.12E-02 | 5.30E-02 | 1.60E-02 | 0.003 | 1.51 | 85.6 | 10.09 | 0.91 |
| Ag 50 nm | adhesion | 2.01E-02 | 2.04E-02 | 2.06E-02 | 2.07E-02 | 2.10E-02 | NA | 0.987 | 102 | 85.7 | 5.19 | 0.97 |

Shown are: Type, silver form; Effect, acute 96 h survival or chronic 24 h byssal adhesion toxicity test; ECx, calculated endpoints; EC50, a parameter of the regression model; SE, standard error of EC50; P, t-test p-value of EC50 as a model parameter; B, hillslope, a parameter of the regression model; Max (%), the constant parameter of the regression model; S, standard error of the estimate; R<sup>2</sup>, squared R of the regression model. All models were significant (F-test, ANOVA, p<0.001); t-test on the predictors EC50 and B failed in byssal adhesion for 50 nm AgNP. All ECx are reported as AUC<sub>h</sub> in mg h L<sup>-1</sup>. NA, cannot be computed.

**Figure 4.**
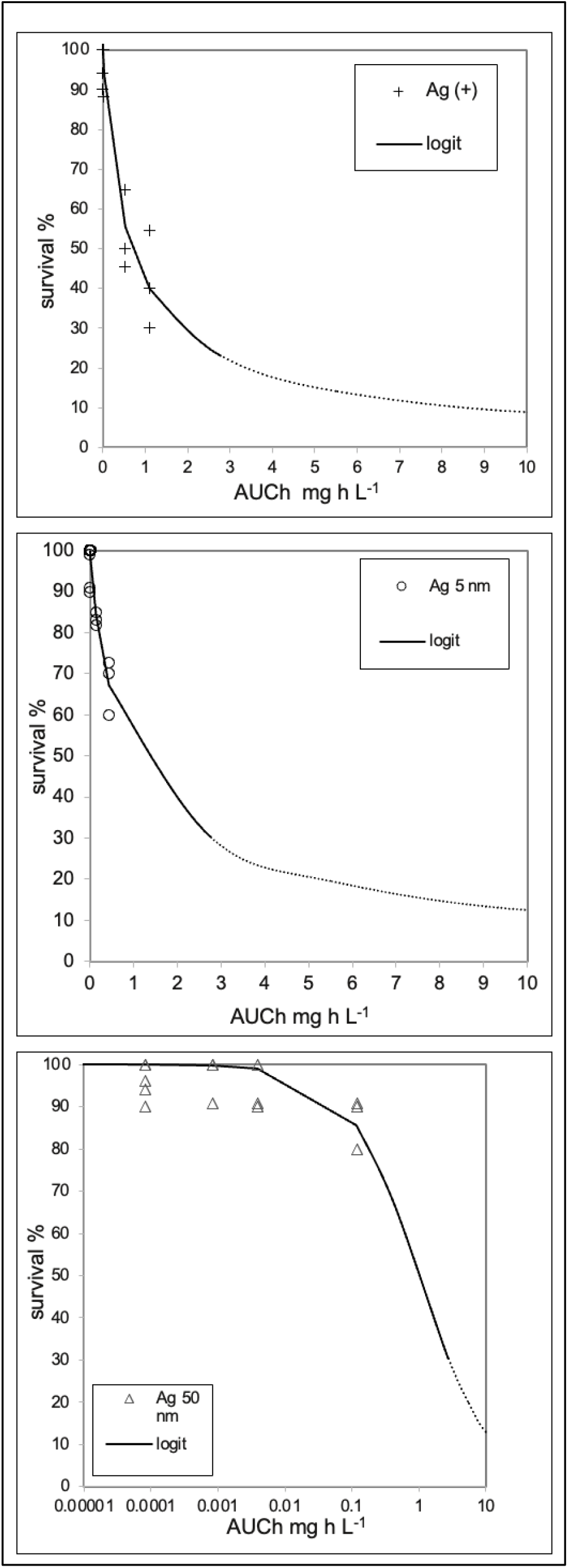
Cumulative toxicity curve for acute toxicity (survival rate). Logistic-regression models using the integrated silver dose as an independent variable. All models proved to be statistically significant by ANOVA. Details of statistics are presented in Table 2.

### 3.3 Chronic toxicity test – byssus adhesion

Byssus adhesion was used as a proxy of chronic toxicity of silver (24 h exposure). Figure 5 shows logistic regression of the effects of the three different silver forms on byssal performance, using the predicted AUC_h_ dose as an independent variable. All models were statistically significant (F test, ANOVA, p < 0.001) although the two parameters EC_50_ and B could not be accurately computed for the 50 nm AgNP (Table 2), most likely due to a lack of observations in the median range. Indeed, the modelling indicated that ionic silver is much more effective on affecting byssal adhesion than 5 nm and 50 nm AgNPs, and in fact, the predicted EC_50_ value is at least a magnitude order lower for ionic silver. To test the statistical significance of such differences, multiple logit regression analysis was performed using the whole byssal adhesion dataset (control not exposed samples were excluded from this analysis, N=49, Chi^2^ < 0.00001) including dummy variables for silver types. This time, the dependent variable adhesion (*sensu strictu*, its odds) can be more accurately predicted from a combination of the independent variables AUC_h_ and the type of silver dummy codes (Supplementary Table 1).

**Figure 5.**
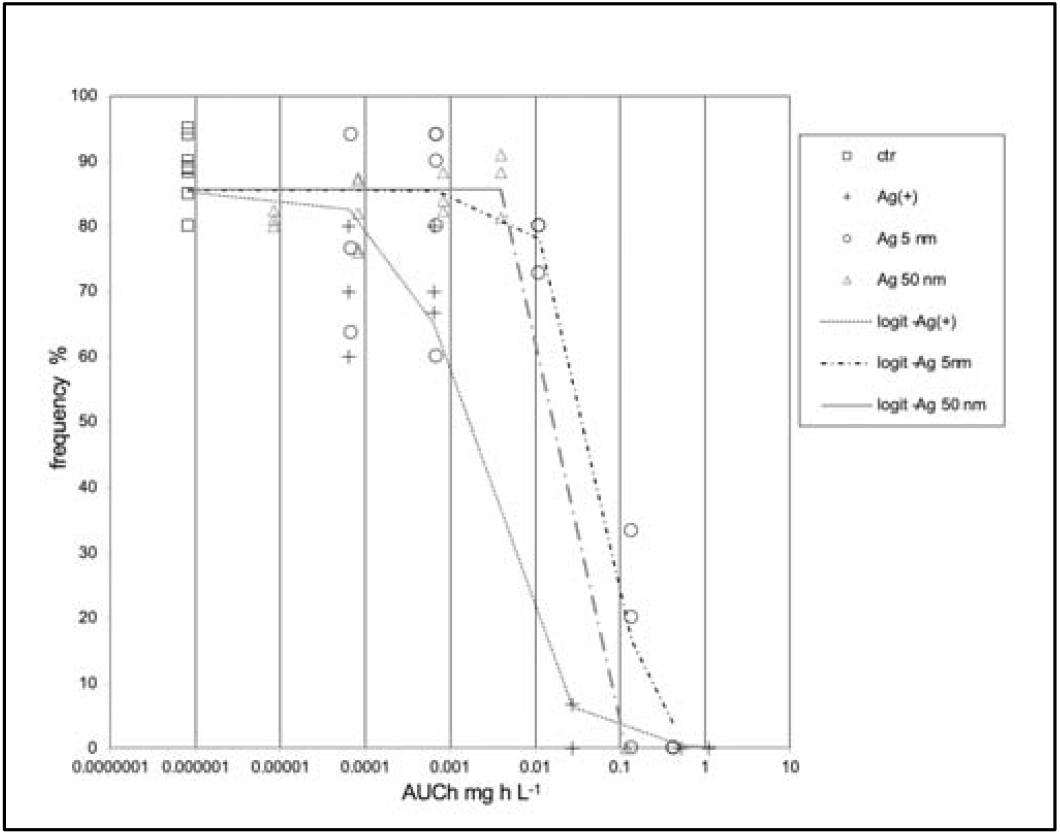
Cumulative toxicity curve for byssus adhesion. Adhesion was used as a proxy of byssus functionality. Shown are logistic-regression models for adhesion obtained using the integrated silver dose as an independent variable. Binomial data were percentualized and logistic regression. The solid line represents an additional model built with all data(see explanation in text). All models proved to be statistically significant by ANOVA. Details of statistics are presented in Table 1.

### 3.4 Silver uptake in mussel body burden

Silver burdens were evaluated in mussel soft tissues after 96 h exposure. Internal silver concentrations could be predicted using a linear regression models with logarithmic transformations (log-log model) with the total (96 h) AUC dose level as the independent variable (Figure 6). The three models obtained displayed similar slopes, suggesting that the uptake is influenced by the silver concentration in a similar fashion. However, the models have different constants, that would indicate different uptake efficiencies, for 5 and 50 nm AgNP, the latter showing the lowest value. To test the significance of such differences, multiple linear regression analysis was performed on the whole uptake dataset using silver type dummy codes as additional independent variables (controls were excluded from the analysis). The internal silver concentration could be predicted with high confidence (N=55 F test, ANOVA, p < 0.001, R^2^ 0.80%) from a linear combination of the independent variables AUC_96h_ (t-test p < 0.001), and dummy codes for silver types, 5 nm AgNP (p = 0.018) and 50 nm AgNP (p < 0.001) (net of a constant). The two dummy variables introduced a negative correction on the log internal concentration respectively of −0.38 and −0.75 fold per each log-AUC_96h_ unit increase of the silver dose of 5 and 50 nm AgNP (Supplementary Table 2).

**Figure 6.**
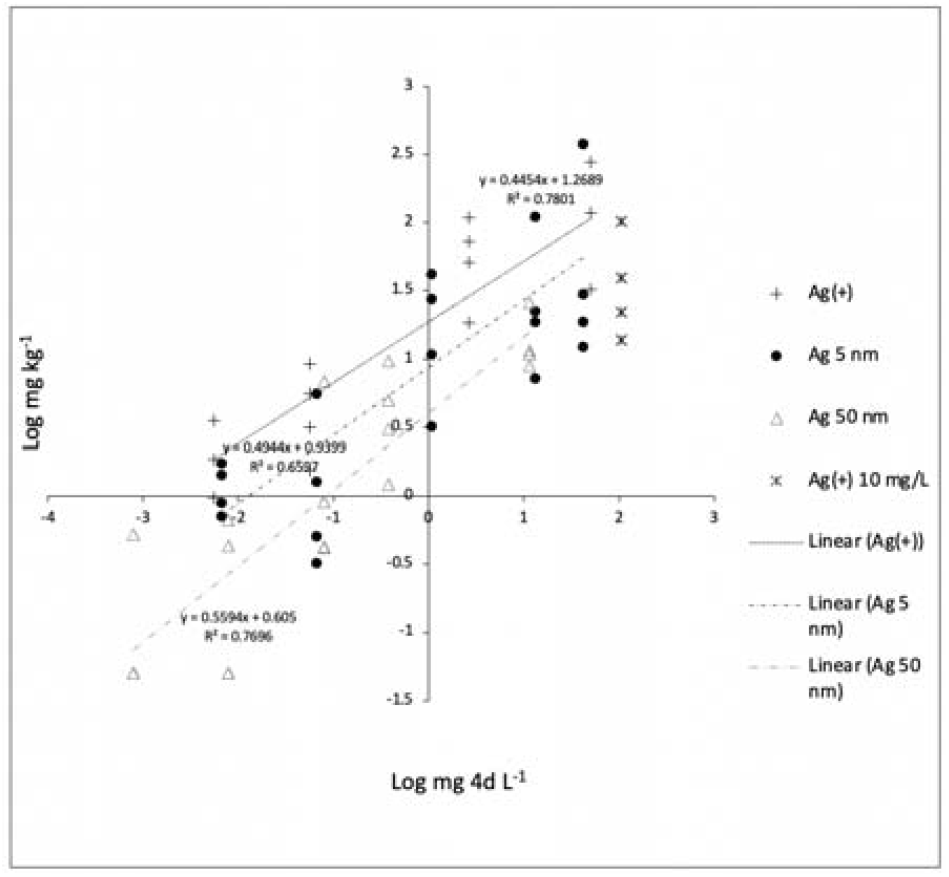
Ag burdens in mussel soft tissues (96 h exposure). Shown are linear regression models (log_10_-log_10_ transformation) using the 96 h integrated silver dose (AUC 4d) as independent variable (data of Ag^+^ at the highest dose level −10 mg L^−1^ nominal concentration-are shown but were excluded from the analysis to avoid any possible bias due to the high mortality rate observed. All models proved to be statistically significant (ANOVA), P value < 0.001. Control not exposed samples were not considered in these analyses.

We also looked at the differential accumulation of silver in the two main bioaccumulating organs, gills and digestive gland after 96 h exposure to the nominal dose of 1 mg L^−1^ (Table 1). Interestingly, ionic silver is almost all found in the gills (95%). This pattern is reversed for 50 nm AgNP showing 59.8 % of silver in the digestive gland. 5 nm AgNP displayed an intermediate pattern, with 85% / 15% repartition of silver in gills or digestive gland, respectively (Figure 7). It’s likely that these patterns are influenced by nanoparticle dissolution.

**Figure 7.**
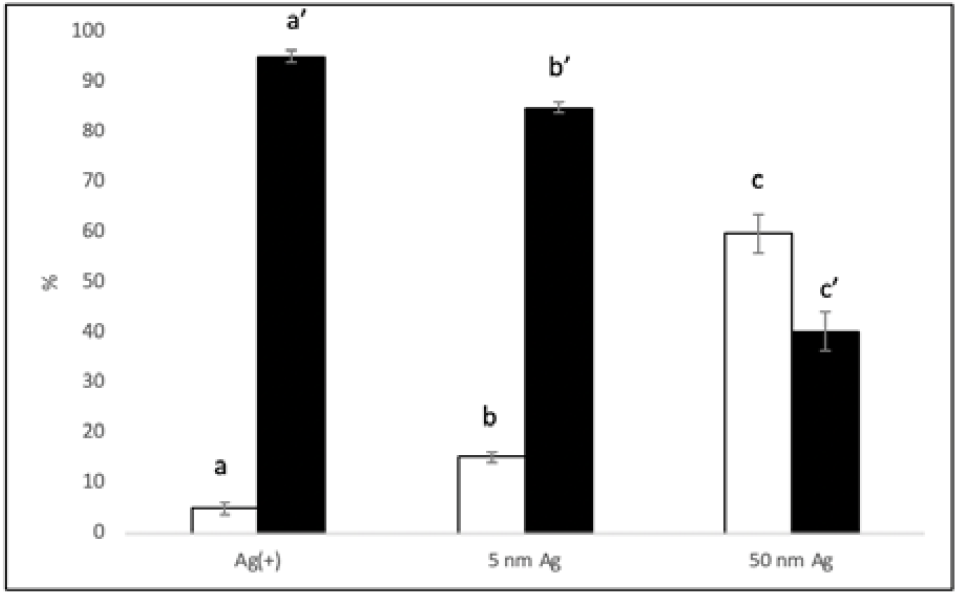
Ag distribution in soft tissues. Silver was evaluated in two main target tissues, gills (solid bars) and digestive gland (open bars) after 96 h exposure to ionic silver (Ag^+^), 5 nm or 50 nm AgNPs. Shown are the average % distributions in tissues found from three independent experiments at 1 mg L^−1^ nominal exposure level. Each experiment used the median value obtained from 8 specimens. Very similar trend was observed also at 0.1 mg L^−1^ (data not shown). A different superscript letter indicates a statistically significant difference (ANOVA, p < 0.0001, N=3)

## Discussion

In this study we provided a framework to address the hazard of AgNPs in the common marine bioindicator species *M. galloprovincialis* Lam, using ionic silver as a reference for toxicity. We embedded two simple and inexpensive tests - 96h acute mortality test and byssus adhesion-in a robust statistical context making extensive use of regression analysis to - first-predict the most accurate exposure levels accounting for silver variation across time and then to test the dependence of toxicity on the actual dose level and the silver form.

A sudden precipitation of silver aggregate with an overall drop of the silver titer was often reported in previous silver nanoparticle exposures in seawater systems (Gomes et al., 2013; Gomes et al., 2014; Ale et al., 2018). Salinity, in fact, is the major factor affecting zeta potential, the electrostatic potential near the particle surface that is a measure of the magnitude of repulsion or attraction between particles (Hochella et al., 2012). In case of ionic silver (provided to the system in form of nitrate) the concentration is expected to remain longer constant due to the formation of complex ions with chloride and hydroxide (Luoma et al., 1995; Santore and Driscoll, 1995; Fabrega et al., 2011). To this end, a persistence model based on the three parameters sigmoidal (logistic) regression function (3PL) was used to fit chemical data-total silver-obtained in time series across 0-24 h, the renewal time in our semi-static exposures. Despite the logistic function is typically used to predict binary data, this model has been successfully and extensively used in toxicology as well as enzymology to describe continuous data by means of parameter iteration minimizing the least square residuals rather than by the maximum likelihood estimation as in canonical logistic models (Haanstra et al., 1985; Jonker et al., 2005). These equations were used to determine the actual silver dose mussels were exposed to as to the Area Under the Curve (AUC_0-24_) rendering in general very good predictions and very low standard errors of the estimation (Table 1). 3PL represents a good model since it relies on a constant that in this case is the maximum silver concentrations in the water column-easily determined from the known amount added to water at time zero; the half-live (τ) of silver concentration in the water column and a hillslope representing the decay rate. As a rule of thumb, τ values associated to AgNPs were often inversely correlated with the initial silver concentration, suggesting the occurrence of concentration dependent aggregation and precipitation processes (Table 1). Moreover, the stability of 5 nm AgNP was much higher than that of larger 50 nm particles. In fact, their τ values, typically, displayed differences higher than one magnitude order. As an example, at 0.1 mg L^−1^ (nominal) level, τ5nm and τ50nm were 1.27 h vs 0.11 h, respectively. At a higher level, 1 mg L^−1^, 0.48 h vs 0.05 h, respectively. At 10 mg L^−1^ level, the precipitation process was instantaneous as argued by the τ values, respectively of 0.07 h and 0.01 h. Half-life for the 5 nm Ag particles could be also computed at 0.01 mg L^−1^ level, rendering a τ equal to 1.19 h. As expected, ionic silver showed the highest persistence in seawater thanks to the formation of complex ions, however, at the highest nominal concentration of 10 mg L^−1^ was clearly fairly above the saturation limit (τ = 0.01 h). In general, at the lowest nominal exposure concentration levels, data could not be fitted, since total Ag was below the detection limit of the technique (LOD, 0.0005 mg L^−1^). In those cases, the AUC was derived using the model parameters obtained for the next higher concentration. Along with AUC_0-24h_ we introduced a linear transformation of it, i.e. AUC per h (AUC_h)_ that would ideally represent a proxy of the actual silver concentration as if this were constant in real scenarios, for example by imagining a continuous input from rivers and discharges to the marine environment.

Once obtained a reliable estimation of the actual exposure levels, we initially used AUC as the unique independent variable to predict silver acute, chronic toxicity as well as silver uptake in mussel soft tissues. For toxicity tests, we again employed the 3PL model that provided accurate predictions, as judged by the overall low standard error of the estimate (Table 2). In a second step, we tested the model improvement by means of multiple regression accounting for the contribution of the different silver form. To this aim, we introduced dummy variables representative of the two AgNP types (ionic silver as to reference), excluding control not-exposed reference samples from the analysis. Despite a slight lower EC50 for ionic silver (Table 2), apical effects, however, could be merely explained by the actual (total) silver concentration in the water column as the two new variables were not significant (see Section 3.2) and did not provide a consistent error reduction to model fitting.

By contrast, a model improvement was determined for byssal adhesion after the introduction of dummy codes for the silver types. Multiple logit regression introduced a statistically significant positive correction on adhesion by a factor 3.66 and 4.65 per each AUC_h_ unit increase, respectively for 5 nm and 50 nm AgNP (as to ionic silver as a reference). This finding poses for an undoubtedly harsher effects of ionic silver over nanoparticles that depends frankly on the silver form.

Silver uptake (bioaccumulation) in mussel body burdens was also measured and modelled as a function of the AUC dose. In this case, we could apply (multiple) linear regression analysis as all conditions for ordinary least square could be satisfied after log-log transformation of the variables. This analysis allowed to infer statistically significantly different uptake kinetics for the three silver forms into mussel soft tissues. AgNPs were, in general, less prone to uptake than ionic silver and thanks to the predicted AUC levels, this difference can be explained through the preferential uptake route of ionic silver vs NPs (Figure 7). The silver amounts found in soft tissues can be correlated with some previous studies, that however are rather heterogeneous. In a study employing a commercial preparation containing AgNPs provided at the nominal level of 0.001 and 0.010 mg L^−1^, after 96 h Ale et al. (2019) found 0.49 and 4.93 mg kg^−1^ (dry weight) in mussel soft tissues, which appear comparable with the range found for 5 nm AgNP at same nominal levels, 0.7-1.7 and 0.3-5.6 mg kg^−1^. Gomes et al. (2014) - who used the same nominal level of 0.010 mg L^−1^ of either ionic silver or 100 nm AgNP in 3–15-day exposure-found a comparable bioaccumulation pattern for gills but significant higher silver amounts were found also in the digestive gland, in particular for silver nitrate. Jemeno-Romero et al., 2017 did not find meaningful silver accumulation pattern in body burdens of mussels exposed for 3 days to either ionic Ag or maltose-stabilized AgNPs (0.00075, 0.075, 0.750 mg L^−1^) of various size, but they justified it for the high mortality rate experienced. They could, however, highlight protein-associated metal deposits in the digestive gland (but not gills, mantle and gonads) by means of auto-metallography. The same research group claimed that dietary exposure is a much better vehicle for silver bioaccumulation (Duroudier et al., 2019; Duroudier et al., 2021).

There are possible explanations accounting for the difference observed between the effects of silver on survival and adhesion. First, the two tests relied on different lengths, 96 h and 24 h, respectively. Then, apical effects such as mortality are a translation of the complex result of molecular initiating and key events as well as their interactions. Previous studies suggested reactive oxygen species formation, protein oxidation (carbonylation), protein sulfhydryl depletion, DNA damage and genotoxicity, that are compatible with an overall electrophilicity of heavy metals ions towards nucleophilic atoms (S, N, O) of biological macromolecules (Gomes et al., 2013; Gomes et al., 2014; Katsumiti et al., 2015; Bouallegui et al., 2018; Duroudier et al., 2021). Proteomic studies indicated also the involvement of moonlight proteins such super oxide dismutase (SOD), glyceraldehyde 3-phosphate dehydrogenase (GAPDH), malate dehydrogenase as well as other molluscan specific features (Duroudier et al., 2019).

For what concerns the effects on byssal adhesion our data demonstrated that adhesive plaques were clearly highly sensitive to silver, in particular the ionic form appeared much more effective than AgNPs. Byssal adhesion relies on a bunch of catecholamines-rich proteins, in particular their modified amino acid 3,4-dihydroxyphenyl-L-alanine (L-DOPA). In *Mytilus* congeners in fact, this modified amino acid is able to establish interactions with natural surfaces and the protein most involved is, of course, mussel foot protein-3 (mfp-3) existing in numerous variants (Zhao et al., 2006). Byssal adhesion relies on the reducing state of L-DOPA residues, hence another cysteine-rich mussel foot protein, mfp-6, acts as an interprotein thiol-mediated redox modulator able to restore the functionality of the corrupted (*sensu* oxidized) adhesive plaque proteins (Yu et al., 2011). This would be a mechanism requiring silver uptake / transport in the mussel foot where adhesive proteins are synthetized, that although we did not verify, it is likely to occur. To this end, reduced thiols (free-thiol oxidation) were reported in mussels tissues exposed to < 50 nm AgNP for just 12h (Bouallegui et al., 2018). We can argue, nonetheless, a second mechanism for byssal adhesion impairment, as functional silky mussel threads establish histidine-mediated intermolecular binding of metals such as zinc at the level of the collagenous stalk (Qin and White, 1998) as well as metal-catecholate complexes at the adhesive plaque matrix protein mfp-1 (Xu, 2013). In this regard, there are clear roles of coordination state and metal types whereas Ag ions - being particularly electrophilic-might further destabilize byssal threads before internalization in a straightforward mechanism from the external. Ionic silver would be immediately available to impair byssal adhesion while AgNPs will require prior dissolution. Despite we did not accurately evaluate the dissolution rate in the water column across time, we gathered preliminary data by ultracentrifugation that suggest a fair amount of soluble silver in the water column. Averaging the three highest nominal exposure levels of 5 nm AgNP, the soluble silver fraction was 14.1 ± 2.9 %, 37.6 ± 5.0 % and 60.3 ± 14.5 % of the total amount found in the water column, respectively at 1, 4 or 24 h. For the 50 nm AgNP, at the highest nominal exposure level, we found a constant 47.5 ± 1.5% soluble vs total silver with no time effects. Some authors reported silver solubilization rates from AgNP in seawater as a function of time. This process seems to be influenced by coating, size and salinity. At 24 h (the renewal time in this work) Katsumiti et al. (2015) counted an overall 9% soluble silver from maltose coated AgNP of various sizes. Schiavo et al., 2017 reported 20% from 5 nm PVP/PEI coated particles but only 1.5% for uncoated 47 nm AgNP; AgNP; Sikder et al., 2018 again for PVP AgNP at 30 ppt salinity reported a ratio of 0.019 % h^−1^, rendering up to 45% soluble silver in 24 h. Gomes et al. (2014) reported a 44% dissolution of < 100 nm AgNP after only 12 h. One of the main pathways of metal ion entry into mussels is passive diffusion across the gill epithelium (George and Pirie, 1980; Scholz, 1980; Carpene and George, 1981; Everaarts, 1990; Soto et al., 1996). Our data on silver uptake dynamics in gills and digestive gland after 96 h exposure indicated the former route is almost exclusive for ionic silver (95%), dominant for 5 nm AgNP (85%) and even for larger 50 nm AgNP (40%) (Figure 7). It should be said that in long-term exposure to ionic silver, in contrast to other heavy metals, silver did not accumulate in the digestive tubules within lysosomes but in the basement membranes and tissue macrophages in some forms associated to sulphur (George et al., 1986) whilst AgNP - in particular larger ones-are more readily available to the digestive tissues (this work; Gomes et al., 2014; Duroudier et al., 2019; Duroudier et al., 2021). Overall, this body of evidence may explain the existence of a double contribution to apical toxicity of AgNP - i.e. via gills and digestive gland by a Trojan Horse effect.

To our knowledge this is the first time in marine nanotoxicology the actual exposure levels, i.e. the AUC, were reconstructed and systematically related to the toxicological outcomes. Unbiased ecotoxicological endpoints were, hence, calculated for acute and chronic toxicity. These values are not affected by the persistence of silver in seawater and are corrected for the actual metal bioavailability / readiness to uptake. This approach allowed us to evaluate the genuine potential of AgNP toxicity and make a reliable comparison with the ionic form. For acute toxicity (survival), we showed comparable LC50 values between ionic and particulate silver. In a previous in vitro study, larger differences were reported ranging between 5 to 21 folds for both hemocytes and gill cells (Katsumiti et al., 2015). In case of chronic toxicity, also our data showed larger disparities between the particulate and ionic silver forms for which we argued mechanistic explanations involving a deferral of silver ion bioavailability dissolving from nanoparticles. Auguste et al. (2018), likewise, essentially attributed to a very low dissolution rate of their 47 nm AgNP the 20-fold change observed between EC50_48h_ values for developmental defects of mussel larvae. They, however, did not consider any sedimentation / precipitation process, therefore, they might have underestimated AgNP toxicity. On the basis of the unbiased ecotoxicological endpoints obtained in our study, it appears clear that in the marine environment the Risk Quotient (i.e. PEC/NOAEL ratio) for nano-silver is too far from providing a level of concern for nano-silver, at least in regard to a chronic risk. Regressed unbiased EC1 values (surrogate NOAELs) obtained in mussels acute or chronic toxicity tests were, in fact, in the range of a few ppb for AgNPs (least value, 2.5 ppb for 5 nm AgNP in chronic test) and down to 20 ppt for ionic silver (Table 2), whereas the Predicted Environmental Concentration (PEC) for nano-silver is as low as 1 pg L^−1^ (Gottschalk et al., 2015; Giese et al., 2018). Even considering a worse case scenarios of 5 ng L^−1^ silver nanoparticle transported by large rivers such as the Rhine (Markus et al., 2016) to an estuarine environment, this would render a negligible RQ value. More attention, however, should be given to the possible contribution of AgNP to the total silver loads from punctual sources, such as discharges and estuaries where the environmental silver concentrations might considerably shift up reaching higher level of concern.

## Acknowledgments

This work was financially supported by the NanoFATE, Project CP-FP 247739 (2010-2014) under the 7th Framework Programme of the European Commission (FP7-NMP-ENV-2009, Theme 4); www.nanofate.eu.

