## Supplemental Table 1 for "Ecotoxicological effects of silver nanoparticles in marine mussels"

Supplementary data 2. Supplementary Table 2. SigmaPlot output for multiple linear regression analysis of silver uptake data

UPTAKE = 1.279 - (0.381 * +RC-"Ag-5nm") - (0.752 * +RC-"Ag-50nm") + (0.490 * AUC4D)

N = 55

R = 0.893	Rsqr = 0.797	Adj Rsqr = 0.785

Standard Error of Estimate = 0.454

**Coefficient	Std. Error	t	P	VIF**

Constant	1.279	0.118	10.869	<0.001

+RC-"Ag-5nm"	-0.381	0.155	-2.455	0.018	1.486

+RC-"Ag-50nm"	-0.752	0.160	-4.700	<0.001	1.579

AUC4D	0.490	0.0427	11.491	<0.001	1.107

Analysis of Variance:

**DF	SS	MS	F	P**

Regression	3	41.218	13.739	66.597	<0.001

Residual	51	10.522	0.206

Total	54	51.740	0.958

**Column	SSIncr	SSMarg**

+RC-"Ag-5nm"	1.600	1.244

+RC-"Ag-50nm"	12.379	4.557

AUC4D	27.240	27.240

The dependent variable UPTAKE can be predicted from a linear combination of the independent variables:

**P**

RC5	0.018

RC50	<0.001

AUC	<0.001

All independent variables appear to contribute to predicting UPTAKE (P < 0.05).

Normality Test (Shapiro-Wilk)	Passed	(P = 0.434)

Constant Variance Test:	Passed	(P = 0.566)

Power of performed test with alpha = 0.050: 1.000

Rc50 and rc5 are dummy codes for 50 nm and 5 nm AgNP respectively; auc is the Area Under the Curve; _cons a constant.
