## Supplemental Table 2 for "Ecotoxicological effects of silver nanoparticles in marine mussels"

Supplementary data 1. Supplementary Table 1. Stata output for multiple regression analysis of byssal adhesion data

```
. fracreg logit adhesion rc50 rc5 auc, vce(robust) or

Iteration 0:   log pseudolikelihood = -32.294159
Iteration 1:   log pseudolikelihood = -23.847199
Iteration 2:   log pseudolikelihood = -19.892712
Iteration 3:   log pseudolikelihood = -18.945781
Iteration 4:   log pseudolikelihood = -18.912442
Iteration 5:   log pseudolikelihood = -18.912296
Iteration 6:   log pseudolikelihood = -18.912296

Fractional logistic regression                  Number of obs     =         49
                                                Wald chi2(3)      =      33.42
                                                Prob > chi2       =     0.0000
Log pseudolikelihood = -18.912296               Pseudo R2         =     0.4426

------------------------------------------------------------------------------
             |               Robust
    adhesion | Odds Ratio   Std. Err.      z    P>|z|     [95% Conf. Interval]
-------------+----------------------------------------------------------------
        rc50 |   3.660316   1.240743     3.83   0.000     1.883586    7.112982
         rc5 |    4.65543   2.048142     3.50   0.000     1.965515    11.02664
         auc |   1.32e-15   1.08e-14    -4.18   0.000     1.40e-22    1.25e-08
       _cons |   1.270657   .4053038     0.75   0.453     .6800162    2.374312
```

Rc50 and rc5 are dummy codes for 50 nm and 5 nm AgNP respectively; auc is the Area Under the Curve; _cons a constant.
